# A digital droplet microarray for measuring tolerance to antibiotics

**DOI:** 10.64898/2026.06.04.730180

**Authors:** Rena Fukuda, Nadia Nikulin, Shenghao Tan, Tobias Dörr, Nate J Cira

**Author notes:** These authors contributed equally to the work (co-first authors) (Rena Fukuda, Nadia Nikulin).

## Abstract

Antibiotic tolerance enables populations of microbes to survive normally lethal antibiotic concentrations, increasing the likelihood of reinfection and facilitating the evolution of resistance. Tolerance measurements typically involve quantifying viable cells after antibiotic exposure. Existing methods range from accessible but low-throughput approaches, such as plate counting, to higher-throughput but semi-quantitative techniques, such as the TDtest. Here, we develop a new system for rapid, precise and high-throughput tolerance measurements. We utilize Surface Patterned Omniphobic Tiles (SPOTs) to discretize cell suspensions into nano-to microliter droplets and estimate the viable cell concentrations following antibiotic exposure from the proportion of empty droplets using Poissonian statistics. We apply the platform to monitor *Klebsiella pneumoniae* tolerance to meropenem over time as a proof of concept. The resulting assay is accessible, compatible with multiple media, and boasts a large dynamic range, sufficient resolution, and rapid handling.

## Introduction

Antibiotic treatment failure is a mounting global health concern.^1^ While the most well-recognized contributor to treatment failure is antibiotic resistance, antibiotic tolerance is increasingly recognized as a mechanism by which microbes evade antibiotic-induced death^2^ and accelerate the evolution of resistance.^3^ Tolerant subpopulations often display phenotypic changes, such as reduced metabolic rate and decreased antibiotic uptake, that enable transient survival at elevated antibiotic concentrations. Tolerance is widespread in clinical pathogens and has been associated with treatment failure and evolution of resistance, but its true impact on clinical outcomes is unclear. This is in part due to diagnostic barriers: while resistance can be quantified using minimum inhibitory concentration (MIC) assays, which are well-established and standardized, no high-throughput, standardized, and truly quantitative test for tolerance exists. The standard practice is timecourse killing experiments using dilution plating and colony counting to measure viable cells; however, this approach is labor- and time-intensive. Higher-throughput systems have been proposed, including the ScanLag system, which quantifies the lag period and growth in colonies as a measure of tolerance.^4^ While automated, throughput is limited due to the need for continuous incubated monitoring of plates. In the TDtest, an adaptation of the Kirby-Bauer disk diffusion assay, nutrients are added post-antibiotic stress to recover growth of persister and tolerant fractions.^5^ While accessible, this method is semi-quantitative and limited in media selection. Thus, there is a need for accessible, media-independent, accurate, absolute tolerance measurements.

Digital assays enable concentration measurements with improved sensitivity and range without complex instrumentation. Several digital assays are employed in microbiology, such as the Most Probable Number (MPN) test. In the MPN test, cell suspensions are partitioned at different volumes in fresh media. If the partition contains live cells, it results in growth. From the fraction of negative (i.e. containing no growth) partitions at certain volumes, one can estimate a cell concentration. The MPN test and other digital measurements provide an absolute quantification of cell viability, unlike optical density and fluorescence measurements, which require calibration curves for each organism. In addition, susceptibility to measurement noise is reduced, since the measurement needs only to discern between a positive and negative instead of monitoring a continuous signal, which may display non-linear behavior. Despite these strengths, the MPN test and other bulk digital assays require many pipetting steps and a large volume of cell suspension for sufficient accuracy and dynamic range, motivating efforts to miniaturize.

A variety of microfluidic systems have been applied to miniaturize digital measurements of cell concentrations,^6,7,8,9,10^ conferring high resolution but limited scalability or accessibility. Droplet microarray systems hold particular promise, as they can rapidly generate many micro-or nanoliter droplets.^11,12^ Here, we utilize our previously developed droplet microarray system, Surface Patterned Omniphobic Tiles (SPOTs)^13^ to generate a high-throughput microfluidic system for tolerance measurements. SPOTs provides parallelized, precise, rapid microdroplet generation of cell suspensions and reagents, without the need for expensive fabrication or imaging equipment. We then apply Poissonian statistics to calculate cell concentration during antibiotic exposure. We employ this assay to monitor tolerance kinetics of the hypertolerant *Klebsiella pneumoniae* TS1 strain and its isogenic mutants with well-defined defects in meropenem tolerance. *K. pneumoniae* is a member of the notorious ESKAPE pathogens, a group singled out for causing particularly worrisome levels of infections that fail treatment.^14^ Despite the increased focus on development of new antibiotics that target resistant strains, difficult-to-treat infections are still rising, particularly for Gram-negative pathogens such as *K. pneumoniae*.^15^ In addition to high levels of resistance, *K. pneumoniae* clinical isolates also often display exceptionally high tolerance levels.^16^ Thus, *K. pneumoniae* was chosen to establish the SPOTs method as a precise high-throughput measure of tolerance to the last-resort antibiotic meropenem.

## Results and Discussion

The SPOTs platform is composed of glass plates that hold droplets, and loaders that deposit liquid onto the plates. Both components have a superomniphobic coating that ensures the platform’s compatibility with a variety of liquids.^13^ The coating on the plate is laser-ablated in circular -philic regions (hereafter referred to as “spots”) of user-defined diameters and positions, where the diameters determine the volume of liquid that each spot partitions. We utilized a customizable loader which is configured to hold four different cell suspensions for enumeration. As the loader is horizontally moved across the plate, droplets are rapidly generated in parallel. We configured the plate and loader to load four cell suspensions, with multiple different volumes per sample (Figure 1A, Table 1). We then loaded growth medium with the redox dye resazurin (a well-established measure of bacterial metabolic activity as a proxy for growth) onto a second plate.^17^ We interfaced the two plates liquid-side to liquid-side (i.e. “sandwiched” the plates)^18, 19, 20^ to combine liquids and prevent evaporation (Figure 1B). The droplet generation process requires less than two minutes per plate (with each plate measuring four samples), demonstrating the rapid liquid handling capabilities of the system.

**Table 1.** Distribution of droplet volumes within proposed layouts.

| Volume ($\mu\text{L}$) | 1 | 0.96 | 0.91 | 0.86 | 0.81 | 0.76 | 0.71 | 0.66 | 0.61 | 0.56 | 0.51 | 0.46 | 0.41 | 0.36 | 0.31 | 0.26 | 0.21 | 0.16 | 0.11 | 0.06 | 0.01 |
| --- | --- | --- | --- | --- | --- | --- | --- | --- | --- | --- | --- | --- | --- | --- | --- | --- | --- | --- | --- | --- | --- |
| Layout 1 | 0 | 0 | 0 | 0 | 0 | 0 | 0 | 0 | 0 | 0 | 0 | 0 | 0 | 0 | 0 | 0 | 0 | 0 | 0 | 0 | 232 |
| Layout 2 | 232 | 0 | 0 | 0 | 0 | 0 | 0 | 0 | 0 | 0 | 0 | 0 | 0 | 0 | 0 | 0 | 0 | 0 | 0 | 0 | 0 |
| Layout 3 | 116 | 0 | 0 | 0 | 0 | 0 | 0 | 0 | 0 | 0 | 0 | 0 | 0 | 0 | 0 | 0 | 0 | 0 | 0 | 0 | 116 |
| Layout 4 | 12 | 11 | 11 | 11 | 11 | 11 | 11 | 11 | 11 | 11 | 11 | 11 | 11 | 11 | 11 | 11 | 11 | 11 | 11 | 11 | 11 |
| Layout 5 (Selected) | 44 | 8 | 8 | 8 | 8 | 8 | 8 | 8 | 8 | 8 | 8 | 8 | 8 | 8 | 8 | 8 | 8 | 8 | 8 | 8 | 36 |

**Figure 1.**
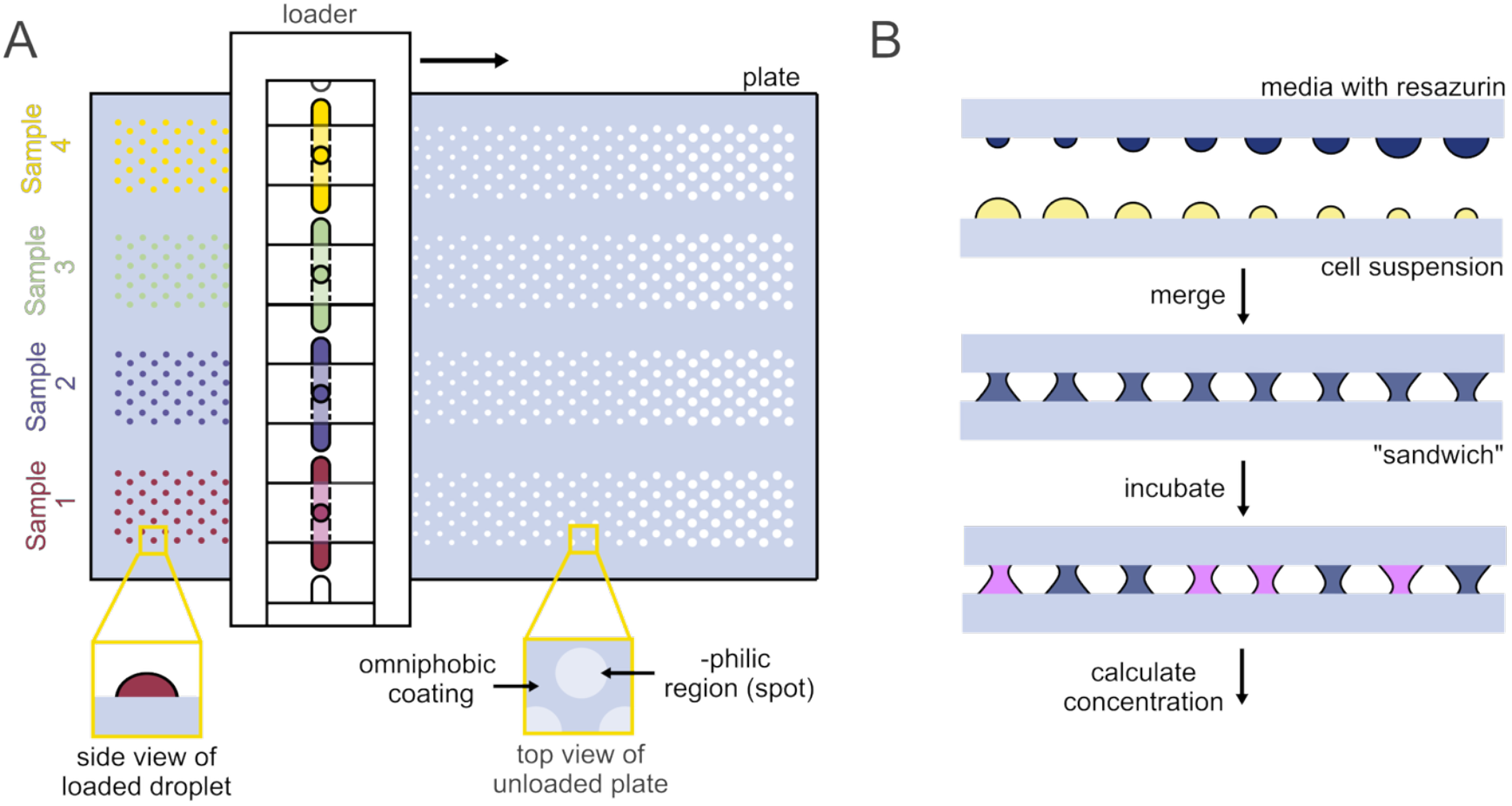
Schematic of assay design. **A.** Samples (four per plate) are loaded using our custom loader, which moves across the plate to generate droplets in -philic regions. **B**. Cell suspensions are interfaced with resazurin in fresh media, then incubated to provide a digital readout.

Following incubation and color change as an indicator of growth, our assay then uses Poissonian statistics to calculate input concentration. Each sample is divided into many discrete droplet volumes. After incubation, due to the metabolic reduction of resazurin, each droplet is visibly positive (pink) or negative (blue) for growth (Figure 1B, 3A). We calculated the input cell concentration using the proportion of negative droplets at each droplet volume, leveraging the multi-volume Poissonian statistics described by *Kreutz et al*.^21^

Tolerance measurements require a large dynamic range. Bacteria often replicate to over 10^9^ cells/mL, and can range from completely tolerant (up to 10^9^ CFU/mL) to highly intolerant (1-10 CFU/mL), necessitating viable cell measurement spanning several orders of magnitude.^16^ The required resolution for tolerance measurements can be relatively coarse, given the large expected differences in tolerance between strains, conditions, and exposure duration (e.g. 10-fold resolution, i.e. able to discriminate between concentration 1X and 10X at 95% confidence or higher). These metrics inform our selection of droplet numbers and droplet volumes, as larger volumes enhance sensitivity to lower concentrations, while smaller droplets increase resolution at higher concentrations.

To devise a suitable assay layout for accurately quantifying tolerance, we considered 5 different schemes, all with the same total number of droplets, which is limited by the plate size and the density at which droplets merge upon interfacing the two plates (Table 1). We computed the dynamic range for each layout (Figure 2A), defined by the lowest and highest concentrations that can be detected with a confidence of 95% or higher.^21^ This corresponds to the concentrations which result in at least one positive droplet or at least one negative droplet in 95% of cases or more, respectively. We also calculated the range of 3- and 5-fold resolution, or the concentration range where one can discriminate between 1X and 3X (or between 1X and 5X) with 95% confidence (1 − *α*) and 95% statistical power (1 − *β*). We then compared these ranges to determine which layout gives the best theoretical performance for tolerance measurements.

**Figure 2.**
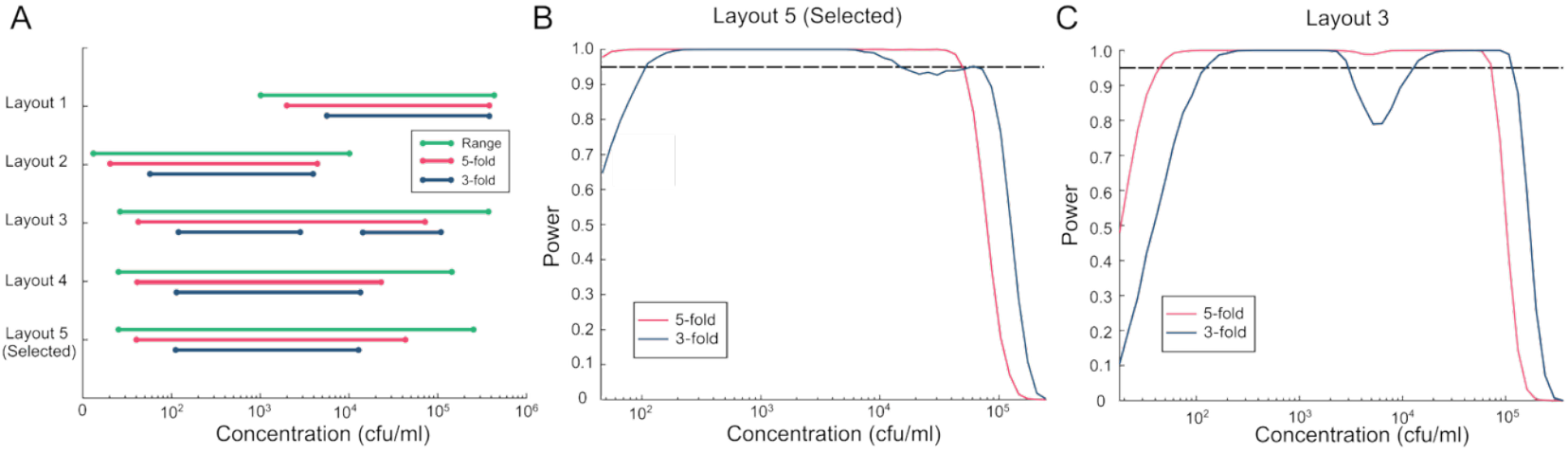
Theoretical accuracy and range of proposed assay configuration and alternative layouts. **A.**Theoretical detectable range, range of 3-fold resolution, and range of 5-fold resolution for layouts proposed in *Table 1*. **B**. Theoretical resolution of current layout (layout 5) and **C**. layout 3 across concentrations. Red and blue lines indicate resolutions of 5- and 3-fold resolution, respectively. Dotted line represents threshold of power (1-β) at 95%, which is the probability of correctly rejecting a null hypothesis – in this case, the probability of correctly distinguishing between two diKerent concentrations (X and 3X for 3-fold, or X and 5X for 5-fold resolution).

We first considered single volume configurations, with Layout 1 containing only the smallest droplets and Layout 2 only the largest. Accordingly, Layout 1 was predicted to effectively cover higher concentrations but fail to detect lowest concentrations, and *vice versa* for Layout 2 (Figure 2A, Supplementary Figure 1). Layout 3 contained a distribution of half large and half small droplets (Figure 2C, Supplementary Figure 1), giving a larger theoretical dynamic range, but with a significant resolution dip in the middle of the range. We next considered Layout 4, a gradient of droplet volumes, which is not expected to have a resolution gap. However, the layout results in a smaller theoretical dynamic range, as the upper end of the dynamic range is set by the number of droplets at the smallest volumes and the lower end of the dynamic range is set by the total volume deposited, which relies more on the number of largest droplets.

Synthesizing all these layouts, our selected layout (Layout 5) is multi-volume, but with more droplets at the highest and lowest volumes. When compared to other single- and multi-volume layouts of the same total droplet number, our proposed layout has a large dynamic range capable of spanning five orders of magnitude, which helps encompass the biologically relevant range of tolerance values. In addition, the current layout maintains 5-fold resolution across three orders of magnitude, exceeding the resolution requirements typically needed for tolerance measurements.

To validate our assay, we next quantified bacterial suspensions of *E. coli* and *K. pneumoniae* TS1 on SPOTs Layout 5 and the predicted cell concentrations with those from actual plate counts (Figure 3B, C). We found a high degree of agreement between plate counts and SPOTs within the detectable range (Figure 3B, C). The concentrations estimated on SPOTs were, on average, slightly higher than those from plate counts, which may reflect discrepancies in growth between liquid medium and agar plates, but were generally within 2.5-fold of plate counts (Figure 3C). A separate experiment quantifying cells in Brain Heart Infusion (BHI), Luria Broth (LB), Muller Hinton Broth (MHB), and wound-mimicking medium (WMM)^22^ also demonstrated high levels of agreement (within 2-fold of plate counts), underscoring the assay’s compatibility with multiple media (Supplementary Figure 2).

**Figure 3.**
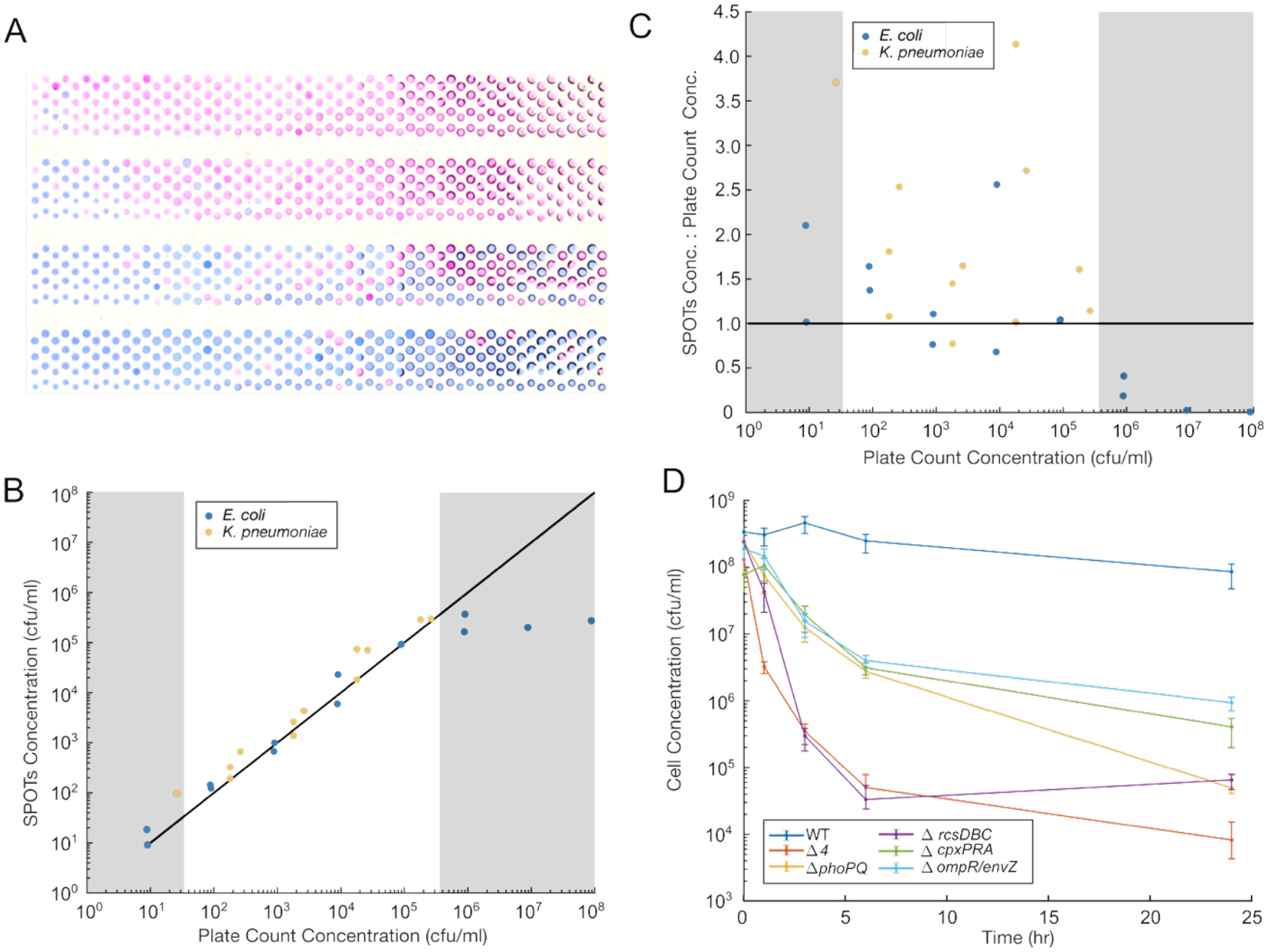
**A.** Sample image of SPOTs plate, with four samples (highest concentration to lowest, from top to bottom). Pink indicates growth; blue indicates no growth. Brightness and contrast were enhanced for viewing. **B**. Predicted versus plate count bacterial concentrations. The predicted values come from the digital SPOTs assay. We assessed agreement for both *E. coli* (blue) and *Klebsiella* (yellow), with the black line representing the perfect agreement. Grayed regions indicate concentrations outside the theoretical dynamic range. **C**. Predicted to known bacterial concentrations ratio, across concentrations. Horizontal line at y=1 indicates line of agreement. **D**. Cell concentrations of six *Klebsiella* TS1 strains during 24 hours of antibiotic exposure, measured on SPOTs. Error bars represent 95% confidence intervals, defined as *CI* = *λe*^±*zσ*^, where *λ* is the cell concentration, *σ* is the standard error, and *Z* is 1.96, the z-score for a two-tailed 95% confidence interval, as further detailed in *Kreutz, et al*.^21^

To demonstrate the application of our platform to tolerance measurements as a proof of principle, we monitored the time course kinetics of the hypertolerant *K. pneumoniae* TS1 strain and its isogenic derivatives exhibiting varying degrees of tolerance^16^ (Figure 3D, Supplementary Figure 3). These mutants have deletions in cell envelope stress signaling systems (alone and in combination), which we have previously shown to be critical mediators of meropenem tolerance. ^16^ As expected from our previously published results, we observed a wide range of tolerance levels within the panel of mutants, from a less than 10-fold viability reduction in the wild-type strain, to greater than 10^4^-fold killing in the *Δ4* strain. We observed rapid killing of *Δ4* and *ΔrcsDBC*, and slower killing of *ΔphoPQ, ΔcpxPRA*, and *ΔompR/envZ*. This is consistent with the understood tolerance phenotypes facilitated by the products of these genes: *ΔompR/envZ, ΔcpxPRA*, and *ΔphoPQ* have a reduced ability to successfully maintain viable spheroplasts in the presence of *β*-lactam antibiotics, a form of tolerance, while also exhibiting recovery defects post-exposure. In contrast, *ΔrcsDBC* mostly exhibits a recovery defect. The *Δ4* mutant is defective in all systems (albeit, it has a deletion in the sensor *rcsF*, while the single mutant is *ΔrcsDBC;* the functional output is lack of Rcs signaling in either case).

Potential adjustments to the assay can further increase throughput, though with certain tradeoffs. Adding resazurin directly into the cell suspensions allows for an open array configuration, removing the sandwiching step. This allows for denser packing of droplets, permitting more droplets per sample and/or more samples loaded per plate which would sharpen resolution. However, this results in more evaporation and weaker signal from smaller droplets. Evaporation may be addressed by encapsulating droplets in oil or submerging plates during incubation, while a higher-sensitivity readout such as fluorescence microscopy can overcome higher noise, though at some cost to readout time and accessibility. If even higher throughput is desired, a kinetic assay allows each sample to be measured with a single droplet. Kinetic assays correlate initial live cell concentration with time to detection.^23^ This can massively increase throughput, but (1) relies on the assumption of a consistent lag time and/or constant growth rate, which are known to vary in tolerant organisms under antibiotic stress, (2) sacrifices the accuracy and noise reduction gained through digitization, and (3) requires time course monitoring as opposed to endpoint, necessitating custom incubated imaging setups.

In summary, we present here a rapid, digital cell-counting assay for measuring antibiotic tolerance. The assay demonstrates high resolution for the application-relevant concentration range, showing improvement over single volume systems. Of note, this platform is versatile and can be applied to other aspects of antimicrobial susceptibility screening, such as across media compositions and strains. The SPOTs platform has demonstrated biocompatibility with a range of microbes, including but not limited to *Staphylococcus pseudintermedius*^24^, *Bacillus subtilis*^25^, *Bifidobacterium pseudocatenulatum, Bacteroides fragilis, Ligilactobacillus animalis*, and *Bifidobacterium longum*.^13^ Future screening efforts may be facilitated by other SPOTs adaptations, enabling a more comprehensive understanding of the factors that influence bacterial tolerance and resistance.

## Materials and Methods

### Preparing SPOTs plates and loaders

Loaders and plates were prepared as detailed in *Shiri et al*^13^; here, we include an abbreviated protocol. The slot and slot-end loader pieces were injection molded using acetyl copolymer (Delrin), then assembled. The SPOTs coating was prepared by combining 300 mg of fumed silica, 10 mL hexane, and 0.3 mL of perfluorodecyl-1H,1H,2H,2H-trichlorosilane (FDTS). The coating was sonicated for 10 minutes. Glass plates and the assembled loaders were sprayed with the superomniphobic coating. The glass plates were baked at 300 °C overnight, while the loaders were baked at 100 °C.

### Enumeration assay

Each cell suspension was first diluted to 10^−3^ and10^−6^ in media with 0.1 mg/mL resazurin. Each dilution was then loaded onto a SPOTs plate. A second plate was loaded with media containing 0.1 mg/mL resazurin. The two plates were interfaced, using a metal spacer to ensure consistent droplet contact and height. These SPOTs plates were incubated overnight at 37 °C in a humidified chamber. In parallel, the cell suspensions were serially diluted, plated on BHI agar, and incubated overnight at 37 °C.

### Time course assay

Six *Klebsiella pneumoniae* TS1 strains (wild type: WT; mutants: *Δ4, ΔphoPQ, ΔrcsDBC, ΔcpxPRA, ΔompR/envZ*) were incubated in BHI media overnight, then diluted 1:100 into fresh BHI. Meropenem (RPI, 10 µg/µl) was introduced at time 0. Then, at 0, 1, 3, 6, and 24 hours, a subsample of each strain was treated with KPC (*K. pneumoniae* carbapenemase purified as outlined in *Cross, et al*)^16^ to degrade the antibiotic, and the cell concentration was quantified with the enumeration assay, using BHI media. At time 0 and 6 hours, samples for each strain were plated on BHI agar at 10-fold dilutions ranging from 10^−1^ to 10^−7^ to serve as a plate count control.

### Plate imaging and analysis

After incubation, each plate was imaged using a digital camera (Canon EOS Rebel T6) in a dark environment, on top of an illuminated LED LightPad. Each image was processed with a custom MATLAB script, which uses user-input points to determine each droplet location, converts the image to HSV, and determines whether each spot is positive or negative, utilizing a threshold value (here a 0.66 hue value). Then, the binary values were input alongside spot volumes into a secondary MATLAB script, which computes concentration using the method adopted from *Kreutz et al*.^21^ The code outputs a concentration value (*λ*) and error (*σ*), which can be normalized by the dilution to produce the final concentration and error.

## Funding

Research reported in this publication was supported by the National Institute of General Medical Sciences (NIGMS) of the National Institutes of Health under award number: R35GM157104. The content is solely the responsibility of the authors and does not necessarily represent the official views of the National Institutes of Health.

## Competing interests

Cornell University has filed a patent application for this technology.

## Notes

### Competing Interest Statement

Cornell University has filed a patent on the SPOTs technology.

