## Supplementary Information for "A digital droplet microarray for measuring tolerance to antibiotics"

### Supplementary Figures

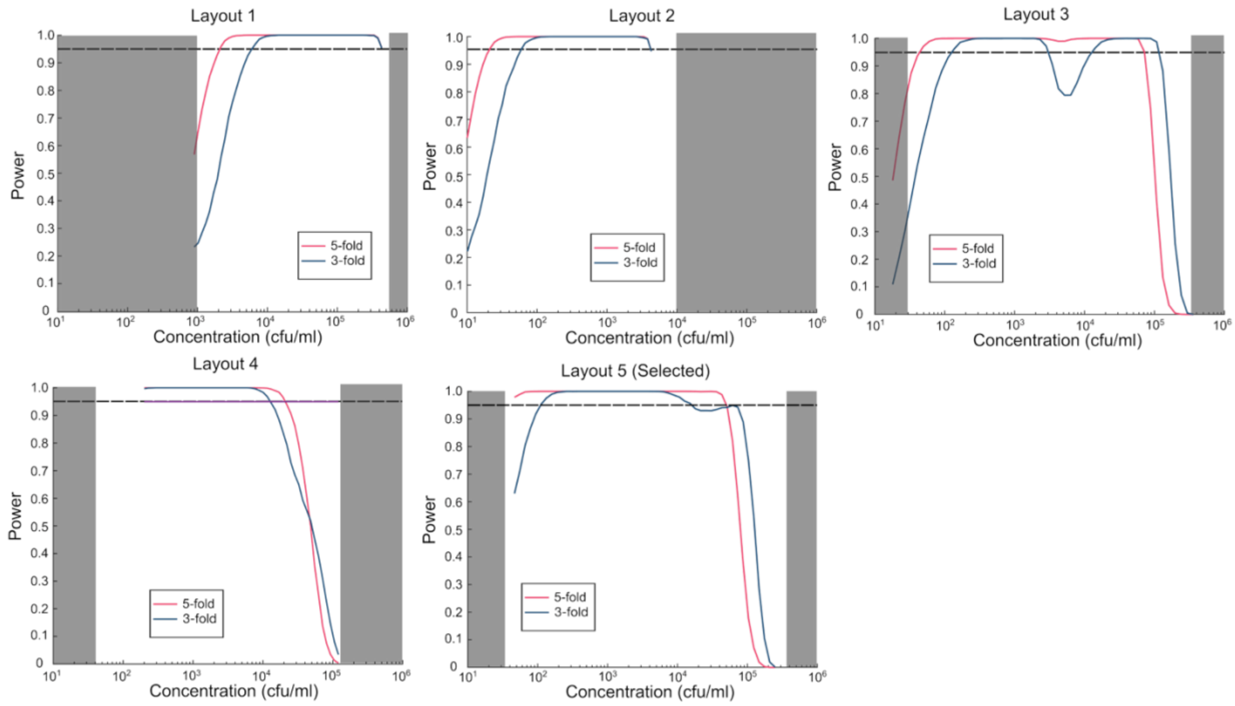


**Supplementary Figure 1.** Resolution of layouts from *Table 1* across concentrations. Red and blue lines indicate resolutions of 5- and 3-fold resolution, respectively. Dotted line represents threshold of power (1-β) at 95%.

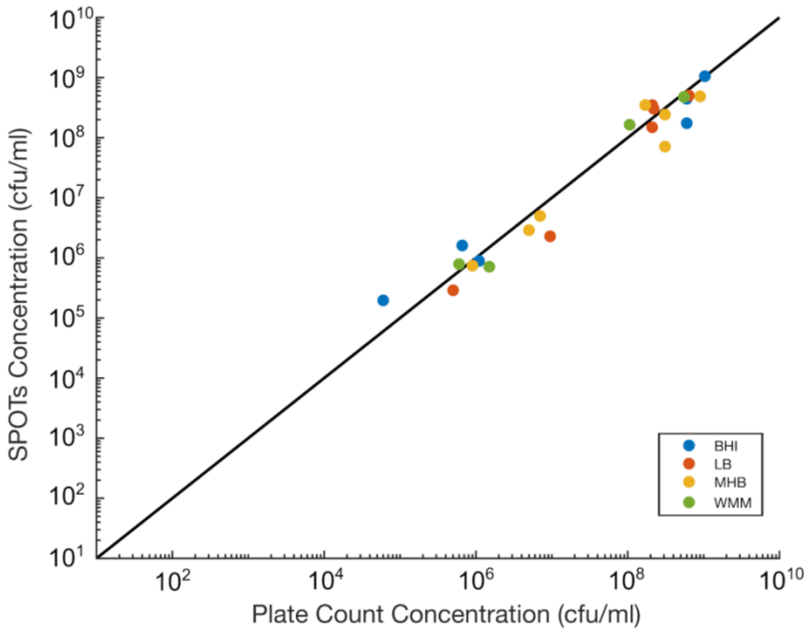


**Supplementary Figure 2.** Concentrations measured using the digital SPOTs assay versus concentrations measured using dilution plating and colony counting. *K. pneumoniae* was grown in Brain Heart Infusion (BHI),

Luria Broth (LB), Muller Hinton Broth (MHB), and wound-mimicking media (WMM). Higher concentration suspensions were diluted to allow for quantification outside dynamic range. Plate counts were conducted on BHI agar. The black line represents the line of agreement.

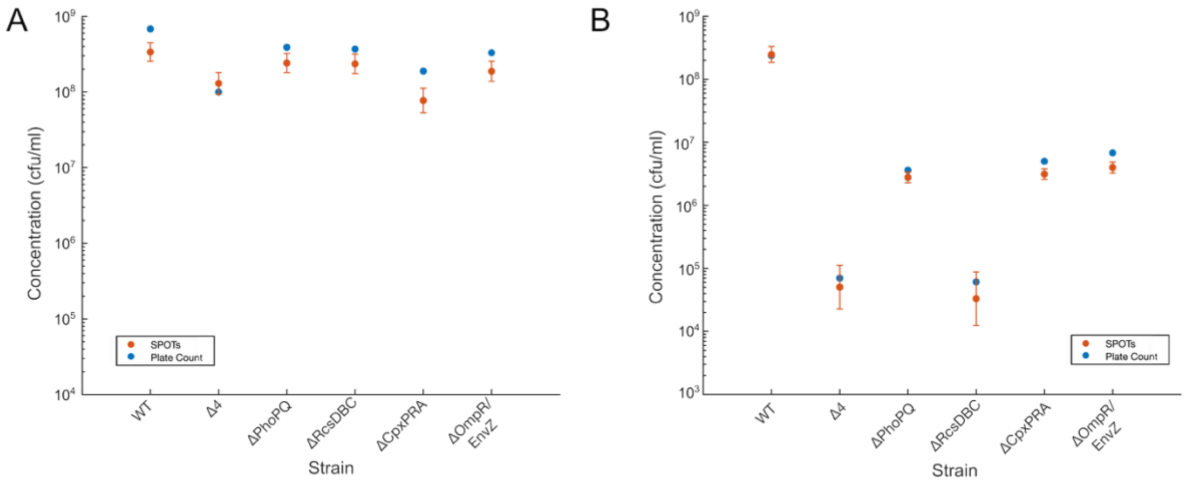


**Supplementary Figure 3.** Cell concentrations measured using SPOTs and plate counts, at time t=0 **(A)** and t=6 h **(B)** during the time course assay for six *K. pneumoniae* TS1 strains.

**Supplementary Video 1.** Protocol of droplet generation and merging. First, one plate is loaded with resazurin using a slot loader. Next, another plate is loaded with cell suspensions (here, colored using food dye for ease of visualization) using a semi-slot loader. The two plates are merged, then incubated before readout.
